# Weighted Network Density Predicts Range of Latent Variable Model Accuracy

**DOI:** 10.1101/343285

**Authors:** Jeremiah B. Palmerston, Qi She, Rosa H. M. Chan

## Abstract

Current experimental techniques impose spatial limits on the number of neuronal units that can be recorded *in-vivo*. To model the neural dynamics utilizing these sampled data, Latent Variable Models (LVMs) have been proposed to study the common unobserved processes within the system that drives neural activities, through an implicit network with hidden states. Yet, relationships between these latent variable models and widely-studied network connectivity measures remained unclear. In this paper, a biologically plausible latent variable model was first fit to neural activity recorded via 2-photon microscopic calcium imaging in the murine primary visual cortex. Graph theoretic measures were then applied to quantify network properties in the recorded sub-regions. Comparison of weighted network measures with LVM prediction accuracy shows some network measures having a strong relationship with LVM prediction accuracy, while other measures do not have a robust relationship with LVM prediction accuracy. Results show LVM will achieve high accuracy in dense networks.

## I. Introduction

Since the late 1950s, analyses of *in-vivo* single-unit recordings in the primary visual cortex have focused on correlating activity of individual neurons with observable features of visual stimuli [8]. However, not all neurons can be confidently classified as maximally responding to a single stimulus feature. Anatomically, it is known that primary sensory regions receive inputs from areas not involved with the stimuli of interest [4].

Imaging techniques have made it possible to record the activity of many neurons simultaneously, *in-vivo*, while an animal is awake [6]. Applying graph theoretic tools on these high-volume simultaneous recordings allows for novel analyses of brain networks. Graph theory can efficiently characterize network types and elucidate important network properties [1]. For example, in the study of sensory perception, this confluence of techniques allows for a change from pairwise associations between individual sensory neurons and sensory stimuli, to network-level measures of sensory representation. Conventional graph-theoretic measures have often evaluated the explicit network where nodes and edges correspond to neuronal units recorded and statistical dependences between these neurons respectively [1]. Yet, the explicit neuronal population dynamics could be modulated by latent variables or implicit network with unobservable nodes and edges, where the implicit activities were not recorded [5].

Therefore, it is plausible that insights into the function of sensory areas can be gained from application of modelling techniques with reduced assumptions on the computational properties of neuronal networks. Recent research on latent variable models provides the mathematical techniques for such models with limited assumptions [7]. One biologically plausible model, rectified latent variable model (RLVM), has been shown to accurately predict the activity of neurons in the rat barrel cortex [11]. Through simulation and application to *in-vivo* 2-photon imaging, the authors show their model accurately describes the activity of the network, and can reveal sensible, interesting, variables driving the activity of the network [11]. A particular strength of this model is that it is able to identify a small number of previously unknown, important factors, driving the activity of many neurons. RLVM has made two important assumptions about the nature of the inputs to the recorded network. First, the latent variables are constrained to non-negativity to simulate neuronal behavior, as there is no neuronal analog of ‘negative activation’. Inhibition is simulated in this model as a negative connection weight from latent variable to cell. Second, the model behaves similar to traditional dimensionality reduction techniques while making no assumption regarding the relationship between latent variables. While such LVMs have been developed to predict the neuronal output from the stimulus and/or other inputs from neurons, the connections between these latent variables and explicit network properties remained unclear. This paper aims to bridge the gap between the modeling frameworks of the explicit network and the implicit network.

## II. Method

### A. Neuronal Recordings

This analysis utilizes 2-photon calcium imaging recordings from the murine primary visual cortex collected by the Allen Brain Institute [2, 3]. The dataset analyzed here includes 488 recordings, with a total cell count of 24,185. Each recording is approximately one hour long, and includes anywhere from 11 to 242 neurons. Each recording is viewed as a separate network with graph-theoretic measures and model application both computed on a per-recording basis. Graph-theoretic measures are normalized by the size of the network in order to compare across recordings. The activity of each neuron is represented as the change in fluorescence at 30Hz divided by the total fluorescence over a 2-second moving-window average (df/f) [15]. All preprocessing is performed by the Allen Institute as detailed in technical documentation; this analysis begins with the df/f trace for each cell [13, 14, 15]. Recordings were obtained from six sub-regions of the murine primary visual cortex. Description of recording locations can be found in [13].

### B. Network Measures

Graph theoretic measures allow for brain networks to be classified and analyzed in previously unexplored ways. In this analysis, six standard, weighted, network measures are employed to efficiently characterize the recorded brain networks. First, the weight matrix of each network is quantified with pairwise Pearson correlation coefficients (R^2^). The absolute value of the correlation coefficient is used to represent each recording as a weighted, undirected graph. Next, graph measures are computed from this weight matrix using Brain Connectivity Toolbox in MATLAB [10]. The *density* of each neuron is computed as average of the linear correlations for each node. Density is used in place of the more common total weight measure, due to the fact that each recording includes a different number of recorded cells. *Clustering coefficient* measures the relationships between groups of three nodes and is computed as a ratio of the sum of weights between such triangles and the largest possible weight (which in this case is 3). The next two network measures are related to distance, where the distance between nodes is represented as the inverse of the correlation coefficient. *Path length* measures the shortest distance between two nodes. Path length includes one value for each pair of nodes in the network, and similar to the density measure above, is represented in the current work as an average for each neuron. *Betweenness* measures the number of shortest paths containing each neuron. This measure results in a single value for each neuron, and is divided by the number of nodes in each network for appropriate comparison across recordings. *Node Centrality* and *Community Structure* are both measures of network modularity [10].

### C. Rectified Latent Variable Model

RLVM will fail to predict neuronal behavior when including neurons with little information (low signal-to-noise ratio). A criterion of signal-to-noise ratio (SNR) greater than one was applied. SNR calculated as a comparison of raw df/f trace to filtered signal as described in [11]. Preliminary analysis showed that as SNR decreases, so does RLVM prediction accuracy. A second preliminary analysis was conducted to evaluate the effect of the number of latent variables on the accuracy of model prediction. The number of latent variables was varied from 5 latent variables, increasing in steps of 10 latent variables, to a maximum number. The maximum number of latent variables was limited for each recording individually to 90% of the number of cells in the network. This preliminary analysis found a plateau of maximum model fit when the number of latent variables reached approximately 80% of the number of cells. A comparison of the model fit with 5 latent variables vs. 80% number of latent variables as cells can be found in Figure 1. 10% of neurons in each region are displayed for clarity, with the subset chosen across the total range of values per region. Each application of RLVM included 5-fold cross-validation as described in [11].

**Figure 1.**
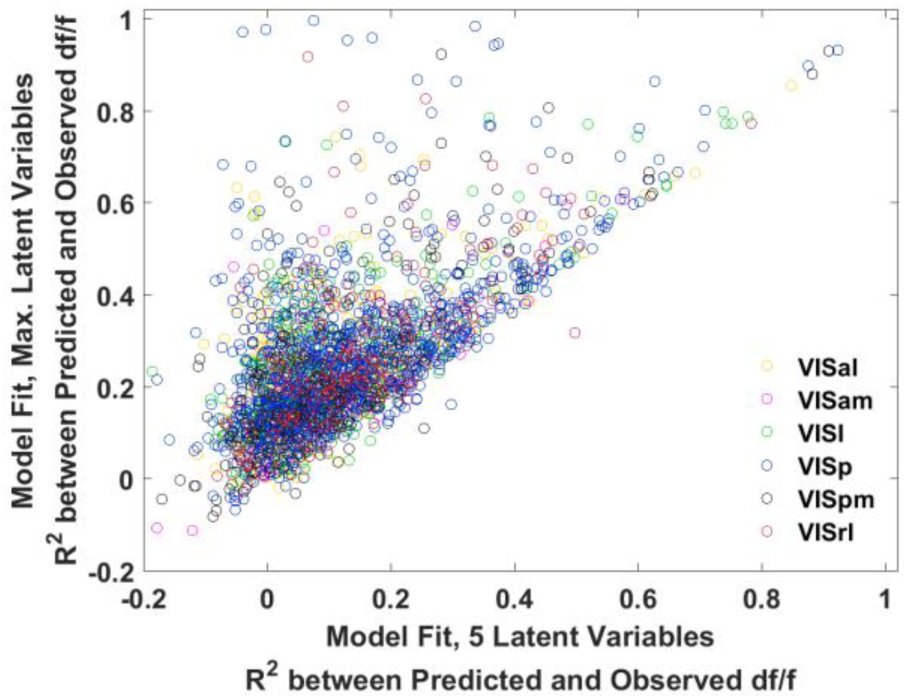
Comparison of Number of Latent Variables: Maximum number of latent variables (80% of number of cells) vs. five latent variables. 10% of analyzed neurons are displayed for clarity. Best viewed in color.

### D. Comparison of Network Measures and RLVM Accuracy

To illustrate the relationship between graph-theoretic measures and the RLVM fit, each network measure was computed to obtain a local value per-neuron and plotted vs. the model fit of the RLVM on a per-neuron basis. Model fit (also referred herein as accuracy) is expressed as the Pearson correlation between predicted and observed neuronal activity. Individual recordings show a range of model fit values across neurons, with most recordings having neurons that were both highly accurately modeled (above 0.6 correlation) and neurons that were not well predicted (close to 0 correlation). Due to the range of model fitting per neuron in each recording, individual-neuron measure comparisons are more appropriate than the recording averages. Pearson correlations (including p-value) between network measures, across all recordings, are displayed in Table 1, while Table 2 relates these network measures with RLVM accuracy per region, also via R^2^. Figure 2 shows node density and corresponding RLVM prediction accuracy for each of the ~24,000 analyzed neurons. There is an observable lower bound in the relationship between density and RLVM accuracy, with R^2^ being highly significant (likely due to large number of neurons). It is also clear that a range of correlations exists between node density and RLVM accuracy. To quantify this observation, quantile regression was applied in 10^th^ percentile increments. Figure 3 plots the regression coefficients between density and RLVM accuracy as well as the intercept values.

**TABLE I.**
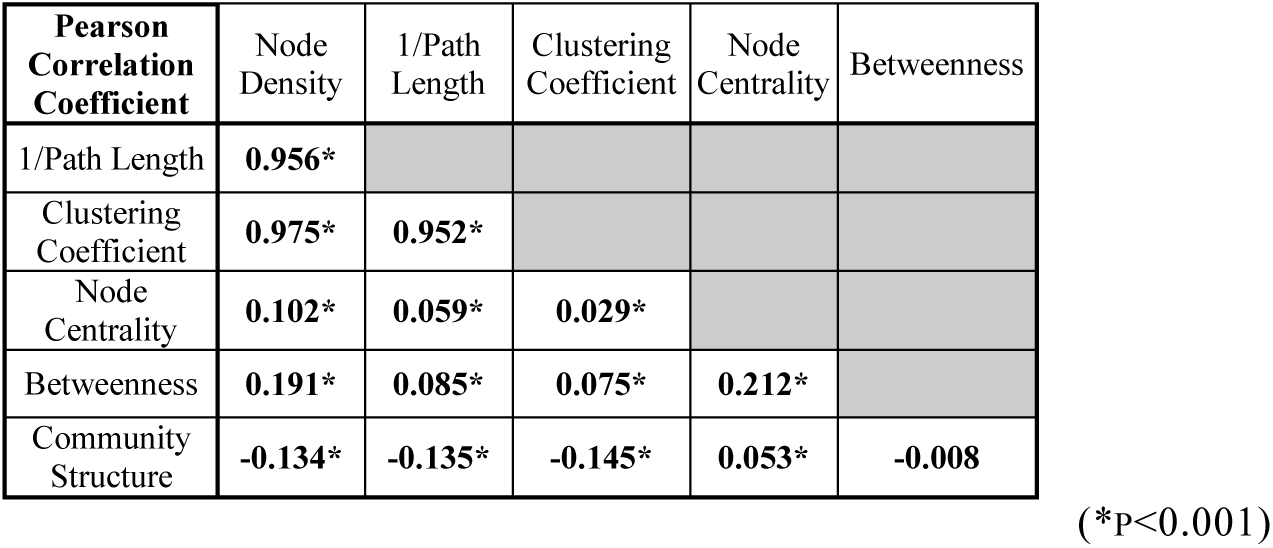
Correlation Between Network Measures

**TABLE II.** Correlation Between Network Measures and RLVM Accuracy

| Pearson Correlation Coefficient | Region |  |  |  |  |  | Average |
| --- | --- | --- | --- | --- | --- | --- | --- |
|  | *VISal* | *VISam* | *VISl* | *VISp* | *VISpm* | *VISrl* |  |
| Node Density | 0.554* | 0.703* | 0.595* | 0.584* | 0.655* | 0.640* | 0.622 |
| Inverse Path Length | 0.536* | 0.635* | 0.609* | 0.582* | 0.599* | 0.650* | 0.602 |
| Clustering Coefficient | 0.366* | 0.534* | 0.528* | 0.497* | 0.512* | 0.551* | 0.498 |
| Node Centrality | 0.319* | 0.308* | 0.252* | 0.250* | 0.330* | 0.223* | 0.280 |
| Betweenness | 0.279* | 0.371* | 0.252* | 0.234* | 0.302* | 0.174* | 0.269 |
| Community Structure | 0.069* | 0.060 | 0.000 | -0.052* | 0.016 | -0.090* | 0.001 |

(\*p < 0.001)

**Figure 2.**
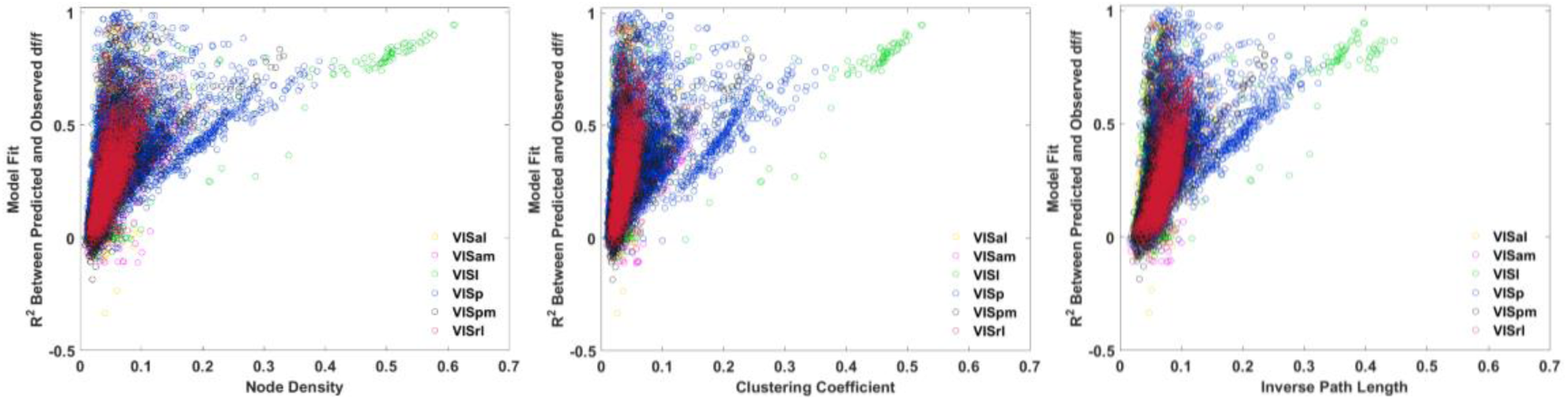
Minimum accuracy of RLVM increases when node density and clustering coefficient increase, and path length decreases, respectively.

## III. Results

Figure 1 illustrates that increasing the number of latent variables typically increases the accuracy of the model prediction. Every sub-region was tested for significance via Student’s t-test. Using a high number of latent variables was found to significantly increase the model fit (all p-values below 0.001). However, although the average RLVM prediction accuracy for each recording increases with increased number of latent variables (as expected), some individual neurons show decreased model fit, suggesting a possible limitation in the model that may provide room for improvement. The points below the diagonal on Figure 1 illustrate this observation.

Table 1 lists observed relationships between network measures; showing density, path length, and clustering coefficient are highly related in definition and observation. Betweenness (which is in this same family of measures depending on node out-weight) of the observed networks does not show a similarly strong relationship with density, path length, or clustering coefficient. Using R^2^ to quantify the weights matrix in this case results in a fully connected matrix, where shortest paths are typically direct connections between neurons. This results in very few paths that include any intermediary neurons and potentially skews the betweenness measure. Node centrality and community structure are slightly more complex measures of networks that have been suggested as important measures for network organization. They are found to have little relationship with the other measures examined here.

Table 2 compares each network measure with the RLVM accuracy. Strong, highly significant, correlations are observed between density, path length, clustering coefficient and RLVM accuracy. This is expected, as highly correlated neurons are likely to share inputs, while the RLVM attempts to predict activity of neurons based on common (hidden) inputs.

Linear regression was applied to fit each network measure, as well as combinations of network measures, to predict accuracy of the RLVM. As expected from the high correlation of density with path length and clustering coefficient, when applying linear regression to predict RLVM accuracy from these network measures, it was found that fitting density to RLVM accuracy achieves a high level of regression fit that is not greatly improved when including more measures in the regression model. When considering application of RLVM to a network, density is an efficient and effective way to quickly estimate how accurately RLVM will predict network activity. It can also be seen from Figure 2 that there is a range of RLVM fit corresponding to similar density measures, with a clear lower bound. This is where quantile regression is useful.

Figure 3 shows the intercepts and regression coefficients for quantile regression between density and RLVM accuracy. Both of these panels illustrate a positive relationship between percentile and regression coefficients. As the percentile increases, so does the coefficient. This means that the lowest quantile represents a lower bound for the strength of relationship between density and RLVM accuracy. Tables 1 and 2 show that density has the strongest relationship with RLVM accuracy, hence density is used as the exemplar herein. The results suggest that node-density can be used to easily estimate a lower bound for expected RLVM prediction accuracy.

**Figure 3.**
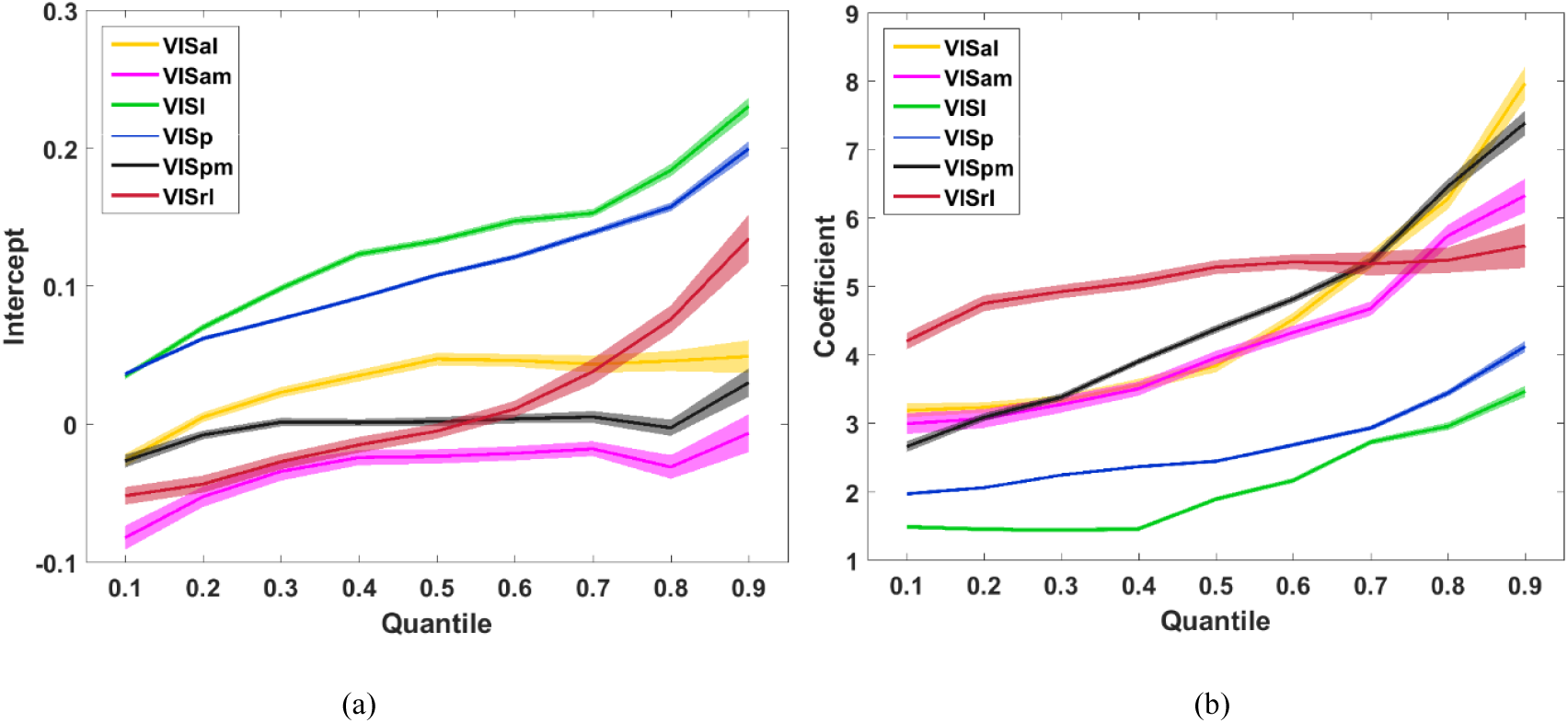
Quantile regression estimates of (a) baseline and (b) weight of node density for RLVM accuracy. The shaded area corresponds to the 95% confidence interval band of the estimate in each region.

## IV. Discussion

Latent variable models attempt to identify the common driving factors of a network and predict the activity of the network nodes based on these previously unknown (possibly unobserved) driving factors. The RLVM allows for limited assumptions to be placed on the nature of the inputs to the network, in order to generate novel observations. This model can be a powerful tool for generating new insights into the functional inputs to brain regions. As the RLVM attempts to identify common factors that drive many nodes in the network, it is reasonable to expect that neurons with similar behavior profiles will be more accurately modeled. This effect is observed, with a high degree of correlation between measures of local connectivity (in this case density, inverse path length, clustering coefficient) and RLVM prediction accuracy.

*In practice,* model selection can be a time-consuming process; this analysis has shown that it is possible to employ simple network measures to identify the applicability of a certain type of model to neuronal data. The method can be extended in three ways 1) via different weight matrix quantification methods 2) with more graph-theoretic measures 3) testing with multiple types of network models. A directed graph quantification can be attained through use of measures such as transfer entropy, Granger causality, or more sophisticated methods of effective connectivity [1]. Investigation of various network measures could elucidate even stronger relationships between graph-theoretic measures and certain classes of network models. The current work suggests these particular network measures may be most suited to estimate a lower bound for prediction accuracy of the RLVM, so that if a network displays high density, short path length, and/or high clustering coefficient, the RLVM is likely to obtain high prediction accuracy per node.

## References

[1] A. Avena-Koenigsberger, B. Misic, O. Sporns, “Communication dynamics in complex brain networks,” Nat. Rev. Neuroscience, vol. 19, pp. 17–33, Jan. 2018.

[2] Allen Institute for Brain Science. Allen Brain Observatory. Available from: http://observatory.brain-map.org/visualcoding, 2016.

[3] Allen Institute for Brain Science. Allen SDK. Available from: https://alleninstitute.github.io/AllenSDK/, 2015.

[4] C. Gilbert, W. Li, “Top-down influences on visual processing”, Nat. Rev. Neurosci., vol. 14, pp.35–363, May 2013.

[5] C. Niell, M. Stryker, “Modulation of visual responses by behavior state in mouse visual cortex”, Neuron, vol. 65, pp. 472–479, Feb. 2010.

[6] C. Stosiek, et al, “In vivo two-photon calcium imaging of neuronal networks”, Proc. Of Nat. Acad. Of Sciences, vol. 100, no. 2, pp.7319–7324, Jun. 2003.

[7] J. Cunningham, B. Yu, “Dimensionality reduction for large-scale neural recordings”, Nat. Neurosci., vol. 17(11), pp.1500–1509, Nov. 2014.

[8] K. Ohki, et al, “Functional imaging with cellular resolution reveals precise micro-architecture in visual cortex”, Nature, vol. 433, pp. 597–603, Feb. 2005.

[9] M. Newman, “Finding community structure in networks using the eigenvectors of matrices”, Phys. Rev., Ed. 74, Sept. 2006.

[10] M. Rubinov, O. Sporns, “Complex network measures of brain connectivity: Uses and interpretations”, NeuroImage, vol. 52, iss. 3, pp.1059–69, Sep. 2010.

[11] M. Whiteway, D. Butts, “Revealing unobserved factors underlying cortical activity with a rectified latent variable model applied to neural population recordings,” J. Neurophysiology, vol. 117(3), pp. 919–936, Mar. 2017.

[12] Q. She, G. Chen, R. Chan, “Evaluating the small-world-ness of a sampled network: functional connectivity of entorhinal-hippocampal circuitry”, *Scienitifc Reports*, article no. 21468, Feb. 2016.

[13] Technical Whitepaper: Overview, Allen Brain Observatory, Allen Institute for Brain Science, v.1, Jun. 2016.

[14] Technical Whitepaper: Phenotypic Characterization of Transgenic Mouse Lines, Allen Brain Observatory, Allen Institute for Brain Science, v.3, Jun. 2017.

[15] Technical Whitepaper: Stimulus Set and Response Analysis, Allen Brain Observatory, Allen Institute for Brain Science, v.1, Jun. 2016.

